# Engineering Cell Fate: The Roles of iPSC Transcription Factors, Chemicals, Barriers and Enhancing Factors in Reprogramming and Transdifferentiation

**DOI:** 10.1101/019455

**Authors:** Behnam Ebrahimi

## Abstract

Direct reprogramming technology has emerged as an outstanding technique for the generation of induced pluripotent stem (iPS) cells and various specialized cells directly from somatic cells of different species. Recent studies dissecting the molecular mechanisms of reprogramming have methodologically improved the quality, ease and efficiency of reprogramming and eliminated the need for genome modifications with integrating viral vectors. With these advancements, direct reprogramming technology has moved closer to clinical application. Here, we provide a comprehensive overview of the cutting-edge findings regarding distinct barriers of reprogramming to pluripotency, strategies to enhance reprogramming efficiency, and chemical reprogramming as one of the non-integrating approaches in iPS cell generation. In addition to direct transdifferentiation, pluripotency factor-induced transdifferentiation or cell activation and signaling directed (CASD) lineage conversion is described as a robust strategy for the generation of both tissue-specific progenitors and clinically relevant cell types. Then, we consider the possibility that a combined method of inhibition of roadblocks (e.g. p53, p21, p57, Mbd3, etc.), and application of enhancing factors in a chemical reprogramming paradigm would be an almost safe, reliable and effective approach in pluripotent reprogramming and transdifferentiation. Furthermore, with respect to the state of native, aberrant, and target gene regulatory networks in reprogrammed cell populations, CellNet is reviewed as a computational platform capable of evaluating the fidelity of reprogramming methods and refining current engineering strategies. Ultimately, we conclude that a faithful, highly efficient and integration-free reprogramming paradigm would provide powerful tools for research studies, drug-based induced regeneration, cell transplantation therapies and other regenerative medicine purposes.

## Introduction

Before 2006, the primary source of pluripotent stem cells was embryonic stem cells (ESCs) derived from blastocysts. By the remarkable breakthroughs and the generation of iPSCs [1, 2], a new window opened to the basic life sciences and regenerative medicine inspired by the previous works of Gurdon [3], Davis [4] and Wilmut [5], which subsequently led to the 2012 Nobel Prize in Physiology or Medicine being jointly awarded to Shinya Yamanaka and John Gurdon. To date, thousands of papers have cited the first induced pluripotent stem (iPS) cells report by Yamanaka in eight years (according to Scopus), demonstrating the importance of pluripotency in the study of early developmental stages, epigenetics, disease modeling, drug screening, cell therapy and regenerative medicine. Direct reprogramming using viral vectors has some concerns such as mutation, genomic alteration and dysregulation in reprogrammed cells, defect in differentiation potential and the risk of tumorigenicity and induction of dysplasia [6-14] caused by insertion of exogenous DNA in the host genome. Interestingly, iPS cells are generated without insertion of exogenous DNA using expression plasmids [15], adenoviral vectors [16], Sendai virus [17, 18], *piggyBac* transposon system [19], permeable reprogramming proteins [20-22], Cre-recombinase excisable viruses [23], excisable *piggyBac* transposition system [24], oriP/EBNA1 (Epstein-Barr nuclear antigen-1)-based episomal vectors [25, 26], synthetic modified mRNAs [22, 27, 28] miRNAs [29], EBNA1/OriP expression plasmid [30], EBNA1-based episomal plasmid vectors [31], small-molecule compounds [32], lipid-mediated transfection of episomal vectors [33], small molecule-aided episomal vectors [34] and episomal vectors in multistage culture system [35]. Such non-integrating methods are reviewed in references [36] and [37-39]. These non-integrating methods could overcome potential safety-related concerns [40]. Moreover, iPS cells can be generated by chemical compounds in a process known as chemical reprogramming using only small molecules and without reprogramming genes [32]. Chemical compounds have various advantages over other methods in iPSC reprogramming [41-44] which will be discussed in a later part of this paper.

Furthermore, the low efficiency and slow kinetics of reprogramming are limitations in iPS cell production in order to clinical applications and regenerative medicine purposes [45-47]. These limitations were significantly resolved by several methods such as inhibition of genetic and epigenetic barriers of reprogramming [48-52] and also overexpression and administration of special enhancing factors [25, 53-58]. Here, reprogramming barriers (roadblocks) and methods/factors that are provided for removing these barriers are discussed as enhancing factors and/or enhancing strategies. Additionally, we will discuss that stem/progenitor cells are more competent resources for efficient reprogramming than terminally differentiated (mature) cells.

Somatic cells can be reprogrammed into different specialized cells using overexpression of lineage specific transcription factor(s) during a process named as direct lineage conversion, direct reprogramming and/or transdifferentiation [59-61]. A versatile transdifferentiation approach that is capable of both generation of terminally differentiated cells, and interestingly tissue specific progenitor cells is cell activation and signaling directed (CASD) lineage conversion [62-64]. Interestingly, the CASD lineage conversion utilizes iPSC transcription factors in conversion of somatic cells (e.g. fibroblasts) into specialized cells and also intermediate progenitor cells without production of pluripotent intermediates [62-64].

As an alternative to reprogramming master genes, we will consider the possibility of application of reprogramming chemicals instead of pluripotent and lineage specific transcription factors in conversion of somatic cells into iPSCs and specialized cells. Theoretically, combining both chemical approach and developed methods of enhancing reprogramming efficiency and kinetics, provides a blueprint toward safe, reliable, and efficient method of pluripotent reprogramming. This proposed combinatorial method cloud be further utilized in transdifferentiation and generation of various cell types. Figuratively, highly efficient chemical reprogramming approaches would provide promises for research studies, clinical applications and induced regeneration. Finally, we have reviewed recent findings about the identity of reprogramming products, the state of native, and target gene regulatory networks (GRNs) in reprogrammed cells, the fidelity of cell fate conversion methods and the capability of CellNet [65, 66], a computational platform for improving reprogramming strategies.

## 1. Reprogramming Barriers, Enhancers and Strategies to Enhance Efficiency and Kinetics

Reprogramming is a time-consuming process and suffers from extremely low efficiency. These features are regarded as limitations for clinical application of iPSCs, although there are hopes and challenges for this purpose [1, 2, 9, 67-69]. Moreover, the species and tissue origins of starting cells have significant effects on reprogramming efficiency and kinetics [70].

Distinct cell types have different requirements for enhancement in efficiency of Oct4, Sox2, Klf4 and c-Myc (OSKM) mediated induced pluripotency [71]. Thus, different strategies should be adopted for efficient reprogramming of distinct cell types. The most widely cell sources that are used in different studies are embryonic, neonatal and adult fibroblasts of mouse and human species. However, this section describes different factors that could act as barriers or enhancers to pluripotent reprogramming regardless to a specific cell source or species (Figure 2).

**Figure 1.**
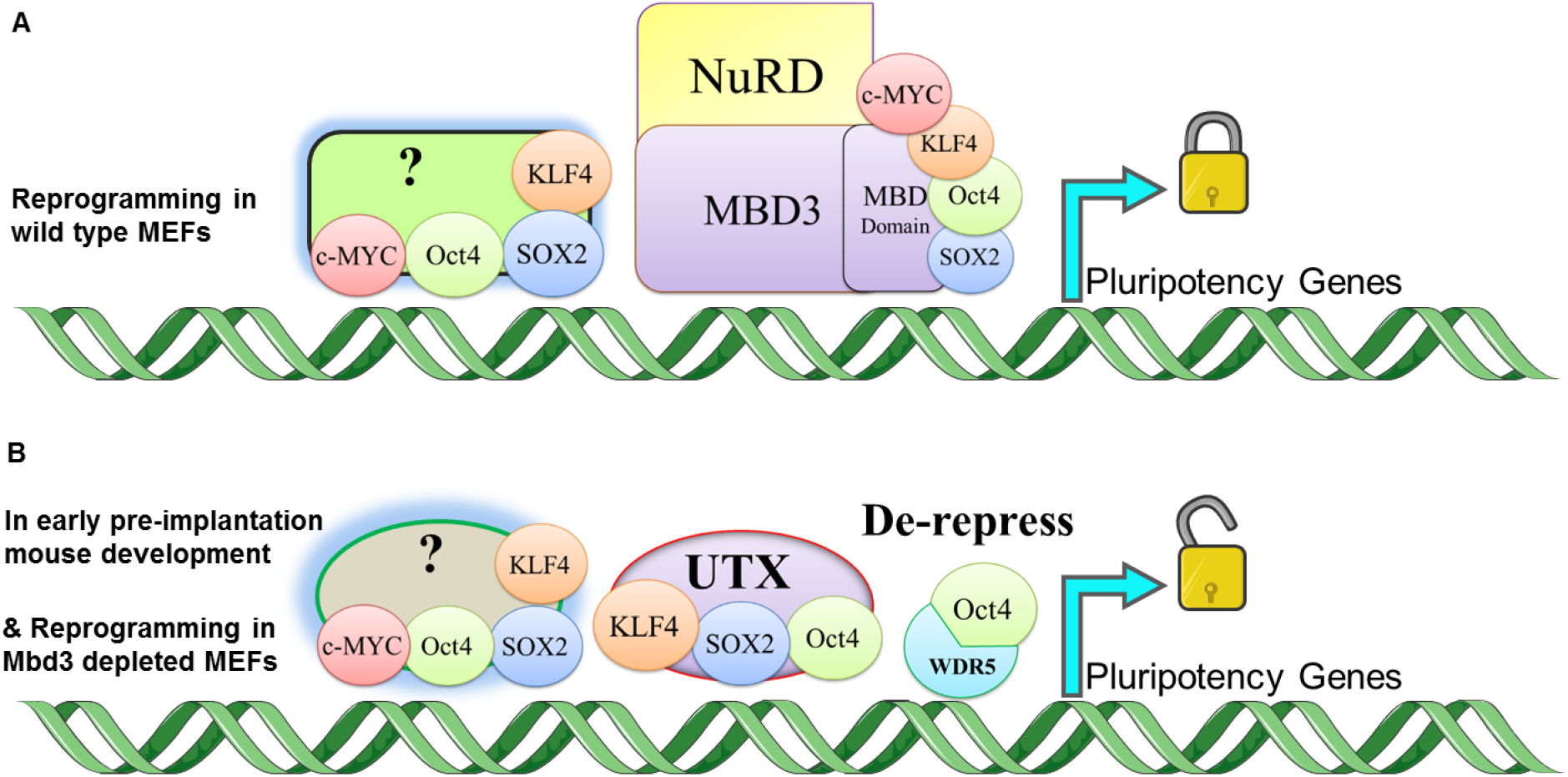
The proposed mechanism by which Mbd3/NuRD complex acts as a reprogramming barrier. (A) During reprogramming, overexpressed OSKM binds to the MBD domain of Mbd3 and other repressive complexes (?) resulting into inhibition of pluripotency-related gene’s expression and reducing reprogramming efficiency. (B) By Mbd3 depletion, reprogramming factors are recruited into downstream targets (e.g. pluripotency-related genes) and consequently, enhance gene expression that leads to highly efficient direct reprogramming. Additionally, WDR5 and UTX are essential in reprogramming of Mbd3 depleted MEFs. It is indicated that Mbd3 is down-regulated in early pre-implantation mouse development and up-regulated in the inner cell mass (ICM) and restricts aberrant specification into the trophoblast lineage [50].

**Figure 2.**
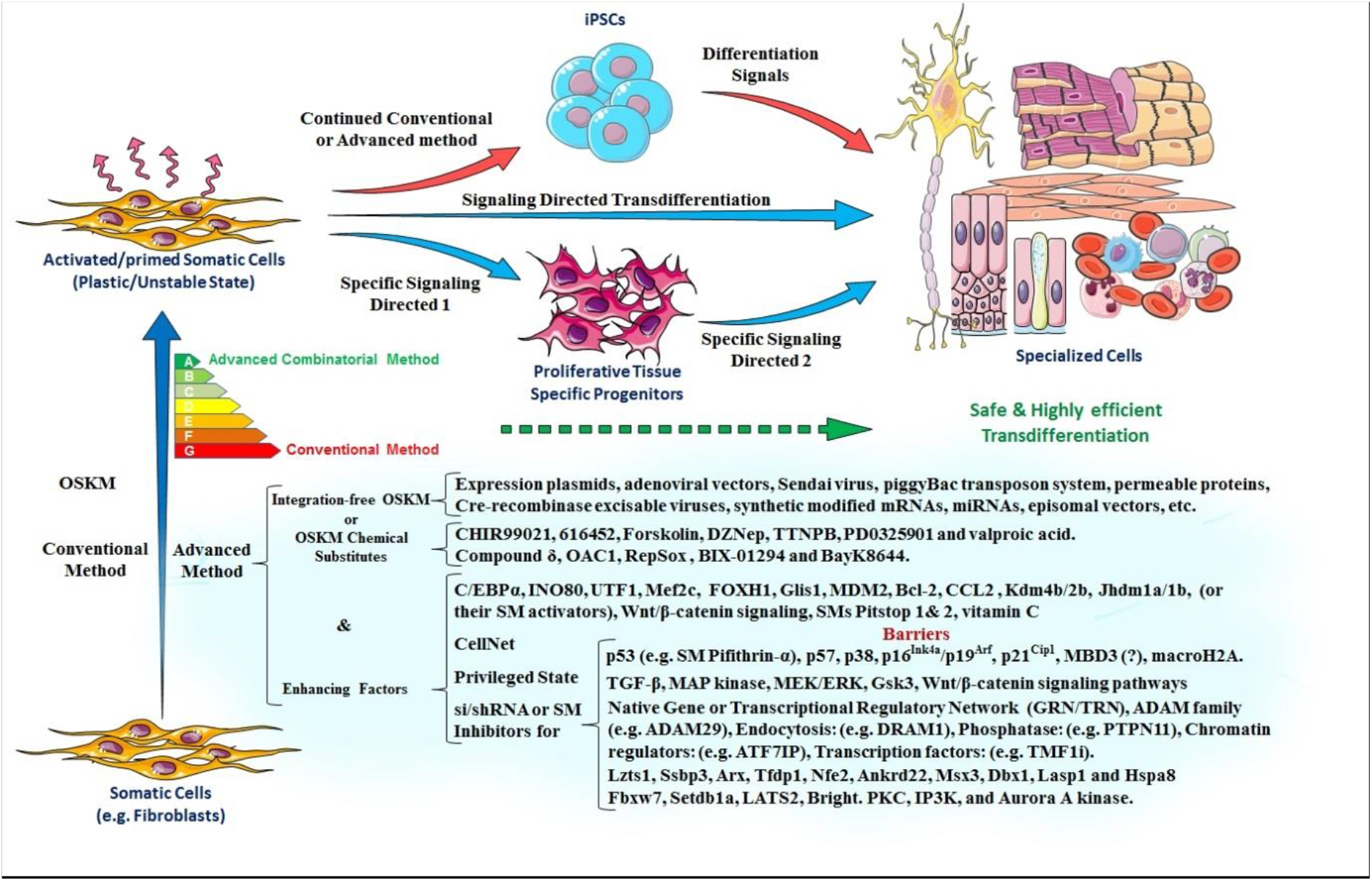
A schematic representation of the topics reviewed in the current paper. Yamanaka conventional method can reprogram somatic cells to induced pluripotent stem cells, and in the subsequent step generated iPSCs can undergo differentiation toward distinct lineages. The CASD lineage conversion system allows for the generation of both differentiated cells directly from activated cells and tissue specific proliferative progenitors. In addition, specific soluble signals can differentiate induced progenitors to the related fates. Interestingly, the mentioned approaches could be done more efficient and safe by using enhancing factors and integration-free OSKM expression or chemical substitutes of reprogramming factors. Enhancing factors include pathways, transcription factors, cytokines, epigenetic modifiers and small molecules that directly are capable of enhancing reprogramming efficiency and kinetics. Moreover, inhibitors of reprogramming barriers are considered as enhancing factors. Various enhancers and roadblocks of the reprogramming are extracted from different reports and listed in the figure regardless of the type of somatic cells and species. As an exception, we included CellNet computational platform in the section of enhancing factors regarding its important role in improving all reprogramming methods. Suggestively, a combined method of removing barriers and administration of enhancers in conjunction with chemical substitutes would powerfully allow for highly efficient reprogramming and transdifferentiation. (SM: Small Molecule).

It is notable that highly efficient and rapid reprogramming methods for the generation of iPS and mature cells are required for certain applications such as cell transplantation therapies, *in vivo* reprogramming, and other regenerative purposes but not essential for cell banking, disease modeling and research studies. Regarding these purposes, here certain advancements on improving reprogramming rate and efficiency will be discussed (Figure 2).

### 1.1. Transcription Factors Affecting Reprogramming Efficiency

Although, forced expression of several known transcription factors (TFs) drives pluripotent reprogramming of somatic cells [1, 2, 67, 68, 72, 73], however, some TFs could act as enhancers or barriers to reprogramming [74-76].

#### 1.1.1. The Gli-like transcription factor Glis1

Glis1 is a Gli-like transcription factor that has been identified as an enhancer of reprogramming and can effectively and specifically promotes iPSC generation from both mouse and human fibroblasts when co-expressed with OSK, in a p53 independent mechanism and by activating several pro-reprogramming pathways. Moreover, Glis1 physically interact with OSK to help the activation of reprogramming target genes. In addition, Glis1 together with OSK can increase ES-like colony formation from human fibroblasts approximately up to 2-fold and 30-fold relative to OSKM and OSK, respectively [75].

#### 1.1.2. The Forkhead box protein H1 (FOXH1)

Yamanaka group have recently demonstrated that transcription factor FOXH1 (forkhead box H1) can facilitate iPSC generation (∼15-fold) when transduced together with OSKM into human adult fibroblasts [76]. Interestingly, FOXH1 facilitates the reprogramming by promoting mesenchymal to epithelial transition (MET) of TRA-1-60^+^ intermediate reprogrammed cells. Moreover, they showed that FOXH1 inhibition during reprogramming can approximately blockages iPSC generation, representative of the important role of this transcription factor in the reprogramming process. Interestingly, it has been revealed that unlike to GLIS1 that facilitates earlier stages, FOXH1 improves reprogramming efficiency by acting at the late stages [75, 76].

#### 1.1.3. The Bright/Arid3A Transcription Factor

Bright/Arid3A is a member of the ARID family of DNA binding transcription factors [77, 78]. Depletion of the Bright/Arid3A transcription factor can confer an increased developmental plasticity and expression of core pluripotency genes to both human and mouse somatic cells, representing this differentiation hallmark as a suppressor of lineage plasticity [79]. Furthermore, Bright/Arid3A has recently identified as a mouse reprogramming barrier by direct binding to the promoter/enhancer regions of *Oct4*, *Sox2*, and *Nanog* and consequently repression of these genes. Popowski and colleagues revealed that depletion of Bright can improve reprogramming efficiency of MEFs for 15 to 40-fold [74]. Moreover, they showed that Bright depletion allows for reprogramming in the absence of Sox2 and Klf4 but not Oct4. Surprisingly, Bright-deficient MEFs can spontaneously form stable iPSCs in leukemia inhibitory factor (LIF) containing medium and without any reprogramming factor [74] that regarding a single performed experiment by these investigators, further demonstrations are needed. In detail, depletion of Bright improves reprogramming efficiency through bypassing senescence, promoting somatic self-renewal, antagonizing differentiation and direct derepression of pluripotency factors [74].

### 1.2. Epithelial-to-Mesenchymal Transition (EMT) and Mesenchymal-to-Epithelial Transition (MET)

Pluripotent reprogramming of somatic cells represents and requires mesenchymal-to-epithelial transition (MET) that is coordinated by suppressing pro-EMT (epithelial-to-mesenchymal transition) signals [80-82]. Interestingly, MET that is promoted by BMP-Smad signaling mediated upregulation of miR-205 and miR-200, is required for and enhances mouse pluripotent reprogramming in initiation phase [83]. In contrast, EMT, the opposite of MET is a developmental process that represents a differentiation process in stem cell and developmental biology; for instance, fibroblasts are products of EMT. Among gene regulatory networks of EMT, transforming growth factor-β (TGF-β) signaling has a critical and dominant role in EMT [80, 81]. It has been shown that TGF-β signaling is a barrier of mouse and human reprogramming and its inhibition can enhance reprogramming [81, 84-86] and also can replace c-Myc or Sox2 in mouse [81, 84, 85]. In addition to discussions above, p53, a known reprogramming roadblock inhibits the process in the early stages by inhibiting MET, and through p21 [87].

Interestingly, iPSC transcription factors progress the reprogramming process by initiating the MET program (e.g. inducing E-cadherin) and shutting down EMT by diminishing intrinsic barriers (e.g. Snail, TGF-β1/TGF-βR2) [81]. However, the presence of TGF-β inhibitor in culture condition removes possible negative effects that could be induced by extracellular milieu [81]. Consequently, EMT and its main regulator, TGF-β signaling, are barriers of mouse and human pluripotent reprogramming and opposing underlying regulatory networks can enhance reprogramming [81, 84-86]. Contrary to this issue, Unternaehrer and colleagues reported that EMT factor SNAI1 (SNAIL) overexpression can paradoxically enhance reprogramming efficiency in human cells and in mouse cells, depending on strain [88]. Thus, although reprogramming efficiency could be improved by preventing EMT and activating MET, but regarding unexpected data more investigation is needed to elucidate the exact roles of MET and EMT during the trajectory of reprogramming.

### 1.3. Barrier Kinases

It has been shown that chemical inhibition of two kinases of Mek/Erk (mitogen activated protein kinase /extracellular signal regulated kinases 1 and 2) and Gsk3 (Glycogen synthase kinase 3) in cooperation with LIF, (2i/LIF) provides an optimum culture condition for the maintenance of ground-state pluripotency in mouse ES cells [89]. It is demonstrated that mitogen-activated protein (MAP) kinase is a reprogramming barrier, and its inhibition can significantly facilitate pluripotent reprogramming of MEFs [71, 90]. Moreover, 2i/LIF promotes significantly maturation step (conversion of pre-iPSC into iPSC) and transition to ground state pluripotency during mouse reprogramming by activating *Nanog* [90, 91]. Likewise, 2i/LIF in combination with a small molecule inhibitor of the protein kinase C (PKC) (Gö6983) can favor induction of ground state pluripotency in human pluripotent stem cells [92]. In addition to above kinases, other kinases including p38, inositol trisphosphate 3-kinase (IP3K), and Aurora A kinase were identified as reprogramming barriers. Furthermore, small molecule-mediated inhibition of these kinases can potently enhance iPSC generation from MEFs [93]. These findings represent inhibitory roles of some kinases (e.g. Aurora A, PKC, MEK and Gsk3) during reprogramming and especially in maturation of iPSCs.

### 1.4. Barrier Signaling Pathways

#### 1.4.1. The p53-p21 Pathway

Different signals can activate p53 that has key roles in the regulation of apoptosis, induction of cell cycle arrest, senescence, and differentiation [94]. It is demonstrated that overexpressed factors (i.e. OSKM), individually or in combination can strongly activate the p53 pathway during reprogramming. Then, this activated pathway impedes reprogramming process and causes a dramatic reduction in reprogramming efficiency [95].

Several groups have shown that p53-p21 pathway is an inhibitor of both human and mouse somatic cell reprogramming and that pluripotent reprogramming could be done more efficient and with accelerated kinetics in the absence of p53 [31, 48, 52, 83, 87, 95-106]. These studies have indicated that direct suppression of p53 signaling pathway can increase reprogramming efficiency of distinct cell types between 10-fold and 100-fold (differs from 100% efficiency). Accordingly, the amounts of increased efficiency are different due to dissimilar culture conditions, reprogramming and knock down systems that were used by the authors [31, 48, 52, 83, 87, 95-106].

It has been revealed that reprogramming efficiency is sensitive to p53 protein dosage and even low levels of its activity can compromise the reprogramming process [95]. Utikal and colleagues showed that secondary OSKM doxycycline-inducible MEFs derived from p53 knockout iPS cells acquire a reprogramming efficiency of about 80% [101] that is striking and indicative of a potent inhibitory role of the p53 pathway during pluripotent reprogramming. However, it is noteworthy that the results elicited from secondary systems or genetically manipulated cells which themselves derived from secondary cells could not be achieved and repeated in direct systems (wild-type cells) and importantly *in situ*. Thus, here, we mostly focus on direct and integration-free systems for induction of highly efficient reprogramming to the aim of cell therapy, *in situ* reprogramming and regenerative medicine purposes rather than genetic manipulation and knockout systems.

Remarkably, p53 restricts reprogramming of mouse and human cells not only by decreasing reprogramming efficiency and kinetics [95, 96, 99-102] but also by elimination of DNA-damaged cells at early stages of reprogramming process via apoptosis [98]. Moreover, p53 depletion allows for efficient reprogramming in the absence of cMyc [99] and cMyc/Klf4 [95]. In addition, *Ink4/Arf* locus which encodes three transcripts of p16^Ink4a^, p19^Arf^ and p15^Ink4b^ tumour suppressors acts as a main barrier of pluripotent reprogramming in both human and mouse somatic cells by activation of p53-p21 pathway, and inhibition of these factors can increase the efficiency of reprogramming [95, 100, 101]. Indeed, inactivation of *Ink4a/Arf*-p53 pathway as a key senescence pathway provides a bottleneck for efficient reprogramming and acquisition of immortality by suppression of *Ink4a/Arf* or p53 removes a key roadblock of the pluripotent reprogramming [101].

Regarding the roles of the p53 pathway in pluripotent reprogramming, it has been shown that MDM2, which negatively regulates p53, can mimic *p53* suppression [99]. In addition, overexpression of the proto-oncogene Bcl-2, an anti-apoptotic protein, can increase the frequency of mouse iPSC formation by fourfold [95]. On the other hand, knocking out or prolonged suppression of p53 reduces the quality of iPS cells and also can lead to genetic instability [99, 107-109]. Moreover, knockout of p53 can cause or increase chromosome end-to-end fusions and chromosomal breaks/fragments in the reprogrammed MEFs compared to wild-type iPSCs [98]. Accordingly, although p53 is a main roadblock of pluripotent reprogramming and decreases the quantity (efficiency and yield), however, increases the quality of reprogrammed cells by induction of apoptosis in suboptimal cells, elimination of these cells and subsequently preventing them from becoming iPS cells. Interestingly, transient suppression of p53 can significantly improve the efficiency of human somatic cell reprogramming using non-integrative plasmids [31, 105, 106]. Furthermore, Rasmussen and colleagues have recently indicated that non-integrative reprogramming approaches in combination with transient p53 inhibition allow for efficient reprogramming without excessive DNA damage due to the presence of low levels of p53 and a reasonable activity of the apoptotic pathway [105]. Remarkably, the main consequence of this strategy is genomic stability of generated iPSCs without any significant effect on apoptosis and DNA damage [105]. Collectively, inhibition of the p53 pathway by small molecules, transiently and in a reversible manner would be a useful tool for enhancing reprogramming efficiency. For instance, small molecule Pifithrin-α as a p53 inhibitor [110] could be used for transient inhibition of the p53 pathway and therefore, enhancing reprogramming efficiency without further genetic instability and malignancies that may arise due to prolonged inhibition of p53 in resultant iPSCs.

#### 1.4.2. Wnt/β-catenin, TGF-β and Hippo signaling pathways

Wnt/β-catenin signaling pathway has differential roles during different stages of direct and cell-fusion mediated mouse reprogramming and temporal modulation of this pathway can considerably increase the efficiency of reprogramming [111-114]. Indeed, repression of Wnt/β-catenin signaling in early stages of Yamanaka reprogramming followed by its normal activity in later stages could significantly enhance the process [112, 113]. This role of the Wnt/β-catenin pathway during mouse reprogramming is almost opposite of its role during cardiomyocyte derivation from the human pluripotent stem cells that needs activation of Wnt/β-catenin in early stages and inhibition of this pathway in the late stages of differentiation [115, 116]. Interestingly, findings suggest that temporally differential behaviors of Wnt/β-catenin pathway are consistent with the activation of MET during establishment of pluripotency and also activation of the EMT during differentiation [80, 113]. Furthermore, Murayama *et al.* have recently reported that inhibition of Wnt signaling can significantly increase the efficiency of mouse EpiSCs conversion to naïve-like pluripotent stem cells in response to LIF, an effect similar to overexpression of E-cadherin [117].

Conversely, it has recently indicated that administration of ascorbic acid together with chemical modulation of two signaling pathways of TGF-β (inhibition) and Wnt/β-catenin (activation) can induce an approximately non-stochastic and highly efficient (80%-95%) OSKM-mediated reprogramming in both somatic and progenitor cells (i.e. MEFs, hepatoblasts and blood progenitors) in a rapid, synchronous and homogeneous manner [71]. In addition, Stadtfeld and colleagues indicated that the only activation of Wnt/β-catenin signaling and the only inhibition TGF-β pathway are sufficient for enhancing reprogramming efficiency of granulocyte monocyte progenitors (GMPs) and hepatoblasts, respectively [71]. Thus, more investigations are needed to elucidate some discrepancies in different reports of the roles of signaling pathways in pluripotent reprogramming.

In addition to the TGF-β signaling [83, 118], that is discussed earlier, another barrier pathway is Hippo signaling that has critical roles in tumor suppression and stem cell function. Interestingly, modulation of this pathway has beneficial effects in anticancer therapeutic strategies and also stimulating tissue repair and regeneration [119]. This pathway has distinct roles in human and mouse iPS cell generation [120, 121]. In detail, Lats2 a serine/threonine protein kinase of the Hippo pathway has been shown to acts as a roadblock in human reprogramming, and its inhibition can improve reprogramming efficiency (∼2.5-fold) [121].

### 1.5. The Ubiquitin-Proteasome System (UPS)

Buckley and colleagues showed that silencing of E3 Ligase Fbxw7 a member of the ubiquitin-proteasome system (UPS) enhances pluripotent reprogramming of MEFs (2-fold) and impedes differentiation of mouse embryonic stem cells (mESCs) through ubiquitination and stabilization of c-Myc [122]. Moreover, *Fbxw7* siRNA can replace exogenous c-Myc expression whereas concomitantly enhances the efficiency [122]. Collectively, recent findings are demonstrating the importance of ubiquitin-proteasome system as a common posttranslational modification in the maintenance of pluripotency in mouse and human ESCs [122, 123] and also pluripotent reprogramming [122] that was less understood.

### 1.6. Insights Gained from Functional Genomics and Genome-wide Studies

In addition to different factors that discussed above, Song and colleagues have recently reported identification of distinct sets of barriers to human pluripotent reprogramming using a novel approach that allows to genome wide screens at an unprecedented scale [118]. They found 956 genes as barriers to reprogramming using their integrative approach [118]. However, among these large numbers of genes several are more effective in hampering reprogramming. These genes are involved in different cellular processes, including, transcription (TTF1, TTF2, TMF1, T), chromatin regulation (ATF7IP, ARID4A, CENPB, MED19), ubiquitination (UBE2D3, UBE2E3, RNF40), dephosphorylation (PTPRJ, PTPRK, PTPN11), endocytosis and vesicular transport (DRAM1, SLC17A5, ARSD), and cell adhesion/motility (ADAM7, ADAM21, ADAM29) [118]. Their results of multiple-and single-gene(s) inhibition of these genes showed significant increases (1.5-fold–15-fold) in reprogramming efficiency [118] comparable with different studies reviewed here.

In line with this approach, Yang and colleagues have recently defined four critical steps in mouse pluripotent reprogramming from initiation to maturation by appropriate markers and applying fluorescence-activated cell sorting (FACS) [103]. Remarkably, using genome-wide RNA interference (RNAi) screen and integrated transcriptome analysis they identified key regulatory genes at each transition step of reprogramming in a stepwise manner. Their findings suggest transition from Thy1^−^ into SSEA1^+^ as a rate-limiting step that expression of pluripotency factors are needed for overcoming this stage [103].

Interestingly, non-differentially expressed genes can act as enhancers (e.g., *Mef2c*, *Utf1* or *Tdgf1*) and barriers (e.g., *Lzts1*, *Ssbp3*, *Arx*, *Tfdp1*, *Nfe2*, *Ankrd22*, *Msx3*, *Dbx1*, *Lasp1* and *Hspa8*) during different steps of reprogramming. For instance, inhibition of *Nfe2* and *Msx3* enhances iPSC generation about 5-fold whereas in the same conditions p53 knockdown enhances 3-fold [103]. Furthermore, inhibition of *Hspa8* and *Lasp1* which act as barriers to reprogramming in maturation step enhance iPSC formation by 8 and 12-fold, respectively [103].

### 1.7. Epigenetic Factors and Epigenetic Modifications Affecting Reprogramming

#### 1.7.1. Histone H3 Lysine 9 (H3K9) Methylation

Histone H3 lysine 9 (H3K9) methylation at core pluripotency genes is one of the epigenetic barriers of mouse pluripotent reprogramming. It is demonstrated that this barrier acts during maturation and stabilization steps and trap reprogramming products in pre-iPSC stage and that removal of this epigenetic mark can terminate the process successfully. [124]. Moreover, it has been revealed that *Setdb1a* histone methyltransferase acts as a roadblock during reprogramming in a BMP dependent manner, and its inhibition can promote conversion of pre-iPSCs into iPSCs with 100% efficiency [124], inconsistent with its function during early human reprogramming [125]. Interestingly, vitamin C can decrease H3K9 methylation through histone demethylases (e.g. *Kdm3a*, *Kdm3b*, *Kdm4c* and *Kdm4b*) at pluripotency loci to further improves reprogramming, an action in opposite of BMP4 [124]. Furthermore, Chen and colleagues showed that *Kdm4b* overexpression can efficiently promote maturation and generation of iPSCs by removing H3K9me3 and H3K9me2, indicative of its rate-limiting activity during complete conversion of pre-iPSCs into iPSCs [124].

#### 1.7.2. Histone H3 Lysine 79 (H3K79) Methylation

H3K79 dimethylation (H3K79Me2) and H3K27me3 mark transcriptionally active and silenced genes respectively. Moreover, lineage specific transcriptional programs could act as barriers to reprogramming to pluripotency (see section 5-2) [126, 127]. Onder *et al.* demonstrated that active H3K79me2 mark acts as a roadblock during reprogramming by hindering repression of lineage-specific programs [125]. Whereas core components of the polycomb repressive complexes 1 and 2, including the histone 3 lysine 27 methyltransferase EZH2, are essential for reprogramming, suppression of histone methyltransferases SUV39H1, YY1 and DOT1L enhances reprogramming efficiency [125]. Furthermore, it has been revealed that small molecule or siRNA inhibition of DOT1L, a histone H3 lysine 79 methyltransferase, can induce a 3-fold to 6-fold increase in efficiency of mouse and human somatic cell reprogramming [125]. Mechanistically, DOT1L inhibition enhances reprogramming by removing active H3K79me2 mark, increasing repressive H3K27me3 mark at fibroblast genes, silencing somatic program and also by a concomitant reverse action on the pluripotency-related genes [125]. In addition, it is demonstrated that the histone variant macroH2A, a differentiation mark that at least in part contributes to the deposition of H3K27me3, is a barrier of reprogramming, and that inhibition of macroH2A.1 and macroH2A.2 isoforms can significantly increase the efficiency of reprogramming [128, 129].

#### 1.7.3. H3K36me2/3 Marks

In line with discussions surrounding the epigenetic modifications, it is demonstrated that methylation at histone H3 lysine 36 (H3K36me2/3) could act as a roadblock during reprogramming and that elimination of the H3K36me2/3 histone marks in some loci is necessary for progression of reprogramming [71, 130]. Moreover, Wang and colleagues revealed that vitamin C enhances removal of these marks via Jumonji histone demethylases, Jhdm1a/1b to potently improves reprogramming efficiency and kinetics even with Oct4 alone [130]. Additionally, overexpression of Jhdm1b (also known as Kdm2b) in conjunction with inhibition of MAP kinase signaling is capable of enhancing efficiency and synchronicity of MEFs reprogramming [71].

Mechanistically, Jhdm1b enhances proliferation of fibroblasts and overcomes cellular senescence by removing H3K36me2/3 marks at *Ink4/Arf* locus and subsequent suppression of this locus that is a known reprogramming barrier [130-132]. Moreover, Jhdm1b removes H3K36me2/3 histone marks from the promoter of microRNA (miRNA) cluster 302/367 to activate these miRNAs as facilitators of reprogramming and subsequently improves the efficiency of reprogramming [130, 133, 134]. These findings delineate, at least in part, the underlying mechanisms that by which Jhdm1b/vitamin C enhance the reprogramming process and also represent the contribution of H3K36 histone modification in both gene activation and suppression during efficient pluripotent reprogramming [130].

#### 1.7.4. Histone Deacetylation

On the role of epigenetic modifications in reprogramming, it has been shown that histone deacetylation and deacetylase enzymes also could act as reprogramming barriers, since histone deacetylase inhibitors can enhance iPSC generation [135-137].

Collectively, chromatin-modifying enzymes play important roles during pluripotent reprogramming, by silencing somatic transcriptional program, activating pluripotency genes and epigenetic remodeling of a differentiated cell state into a pluripotent state. By attaining mechanistic insight into the specific activities of epigenetic enzymes as facilitators or barriers and also by modulating their activities or expression, highly efficient reprogramming could be achieved with reduced kinetics and even with fewer factors.

#### 1.7.5. The MBD3/NuRD Complex

Reprogramming is principally an epigenetic process and chromatin modifiers have critical roles in genome remodeling. In addition to the current discussion, the epigenetic changes that facilitate iPS cell reprogramming are reviewed by Buganim and colleagues [138] and elsewhere [125, 139, 140]. Efficiency of direct reprogramming into specialized cells (e.g. induced neurons >20% [141] and induced cardiomyocytes=20% [142],) is reported higher than iPS cell production (<0.1% [1]) which may suggest epigenetic barriers as an obstacle in reprogramming to pluripotency. Fidalgoa *et al*. showed that pluripotency-related transcription factor Zfp281 directly recruits NuRD repressor complex to the *Nanog* locus and subsequently restrict *Nanog* reactivation and inhibits iPSC formation, representing an inhibitory role for NuRD complex in pluripotent reprogramming [143]. Conformingly, Luo and colleagues showed that methyl-CpG-binding domain protein 3 (Mbd3), a subunit of the NuRD, impairs pluripotent reprogramming of MEFs and that its inhibition can improve reprogramming efficiency even in the absence of c-Myc or Sox2 [49]. They showed that Mbd3 as a roadblock of reprogramming, downregulates pluripotency specific genes (*Nanog*, *Oct4* and *Sox2*) and its depletion can lead to upregulation of these genes and improving reprogramming efficiency [49].

Confirming these findings, Rais and colleagues showed that Mbd3 is a key molecular barrier preventing deterministic (with 100% efficiency) induction of ground-state pluripotency [50]. These authors showed that inhibition of Mbd3 increases the efficiency of mouse epiblast stem cells (EpiSCs) reversion into naïve pluripotent cells up to 80% [50]. Moreover, genetically Mbd3 depleted MEFs under optimized reprogramming conditions (2i/LIF) were reprogrammed into iPS cells with 100% efficiency by day 8 in comparison with 20% reprogramming efficiency in wild-type MEFs in the same conditions [50]. Remarkably, adult progenitor cells, including, common myeloid progenitor cells (CMP), hematopoietic stem cells (HSC), neural precursor cells (NPC) and also terminally differentiated cells (monocytes, Mature B and T cells) were reprogrammed with 100% efficiency under these conditions [50]. However, the roles of GSK3β and MEK signal inhibition and LIF are undeniable in Rais reprogramming cocktail. Surprisingly, in an apparent disagreement with the findings of Fidalgoa *et al*. [143], Luo *et al.* [49] and Rais *et al.* [50] that gave emphasis to the inhibitory role of Mbd3/NuRD complex during pluripotent reprogramming, dos Santos and colleagues have recently reported generation of reprogramming intermediates or preiPSCs from Mbd3^−/−^ neural stem cell (NSC) line by cMyc, Klf4, and Oct4 (MKO) and then serum plus LIF (S+LIF) conditions, but less efficient and with delayed kinetics [54]. However, the inhibitory role of serum in the dos Santos *et al.* reprogramming medium is undeniable as demonstrated earlier [124]. Furthermore, Dr. Silva’s group showed that Mbd3 depletion strongly impairs the conversion of NSCs into preiPSCs in the initiation phase of reprogramming and also strongly reduces efficiency of conversion to naïve pluripotency [54]. They demonstrated Mbd3 requirement in the initiation phase, but not in the establishment of pluripotency in NSC reprogramming [54], on the contrary of Rais *et al.* [50] that showed inhibitory activity of Mbd3 before establishment of pluripotency in early stages. Moreover, Mbd3^−/−^ iPS cells showed slower proliferation and impaired embryoid body differentiation [54]. Strikingly, results of dos Santos and colleagues showed that Mbd3 depletion impairs reversion of EpiSCs to naïve pluripotency [54]. Their results showed that depletion of MBD3/NuRD cannot enhance reprogramming efficiency of MEFs and moreover, its overexpression has no positive or negative effect on the efficiency, depending on the reprogramming context [54]. These findings are in apparent disagreement with the results of Rais *et al.* [50], Luo *et al.* [49] and Fidalgoa *et al*. (including Dr. Jose Silva) [143]. Additionally, some recent studies also did not find an inhibitory role for Mbd3 during mouse and human reprogramming using genome-wide RNAi screen [103, 118] and selective RNAi screen [125]. Regarding these discrepancies, Silva and colleagues raised a concern that there are some methodological issues in Rais *et al*. that negatively affect accurate interpretation of the results [144]. Thence, Hanna and colleagues [145] in disagreement with Bertone *et al*. [144] confirmed the inhibitory role of Mbd3/NuRD pathway in the maintenance and induction of pluripotency by providing new data [145]. Most recently, Wernig and colleagues independently confirmed the validity and authenticity of Rais *et al*. method and deterministic reprogramming [146]. This topic has recently attracted much attention and become a controversial issue in the field. Thus, further investigations are needed to elucidate the basis of contradicting and also striking findings of these groups.

##### 1.7.5.1. Molecular Mechanisms Underlying Mbd3 Interactions

Hanna and colleagues reported that before OSKM overexpression, Mbd3 and Chd4 (NuRD components) are not localized to pluripotency factor target genes; however, after OSKM induction, Klf4, Oct4, Sox2 and Esrrb target genes become enriched for Mbd3 and Chd4 binding. Interestingly, these target genes become significantly up-regulated upon depletion of Mbd3, which indicates that the Mbd3/NuRD complex is a repressor of reprogramming [50]. After OSKM overexpression, Mbd3 binding shows an 8-fold increase that resembles a brake in somatic cells resisting against conversion into pluripotency (Figure 1a). It is indicated that depletion of Mbd3 increases Oct4 binding, H3K4me3 and H3K27ac (derepression marks) and decrease H3K27me3 (repressive chromatin marks) during reprogramming [49, 50]. In addition to OSKM expression, Utx and Wdr5 are also essential for reprogramming in Mbd3-depleted cells (Figure 1b) [50].

During reprogramming, exogenous OSKM proteins bind to the MBD domain of Mbd3 and a direct interaction of Mbd3/NuRD and Chd4/NuRD with overexpressed OSKM recruits these complexes into their own promoters and downstream targets of Klf4, Oct4, Sox2 and Esrrb that consequently cause repression of pluripotency genes. NuRD and OSKM could not assemble a repression complex in the absence of Mbd3 and subsequently OSKM and downstream targets be activated under continued expression of exogenous OSKM (Figure 1b). Accordingly, the Hanna group described this process as a ‘gas and brakes’ paradigm. This paradigm shows a biphasic role for OSKM that both drives reprogramming forward while also inhibiting reprogramming by binding to Mbd3 [50].

Surprisingly, dos Santos and colleagues have indicated that overexpression of Mbd3 not only is not a barrier but also in conjunction with overexpressed Nanog can increase reprogramming kinetics and efficiency in MEF-derived preiPSCs [54]. They demonstrated that overexpressed Nanog interacts with NuRD and in the presence of overexpressed Mbd3, the Mbd3/NuRD complex can enhance reprogramming efficiency of MEF-derived preiPSCs and EpiSCs [54]. Contracting this finding, Mbd3 was reported as a barrier of reprogramming in the late stages by silencing *Nanog* [49].

### 1.8. The CCAAT/Enhancer Binding Protein-α (C/EBPα)

In attempting to increase the efficiency of reprogramming Dr. Graf’s group has recently reported an efficient method for reprogramming of mouse committed B-cell precursors (B cells) into iPS cells [53]. Interestingly, they found that transient expression of CCAAT/enhancer binding protein-α (C/EBPα) for an optimized 18-hour followed by OSKM expression can induce iPS cell reprogramming in B cells with 95% efficiency and emergence of Oct^+^ cells in only 2 days. In detail, direct binding of the overexpressed C/EBPα to methylcytosine dioxygenase *Tet2* leads to up-regulation of this gene and then binding of TET2 to the regulatory regions of the pluripotency genes. This action leads to oxidation of 5-methylcytosine (5mC) residues into 5-hydroxymethylcytosine (5hmC) and consequently this modification could induce demethylation, chromatin remodeling and gene’s de-repression and makes pluripotency gene promoters more accessible for the Oct4 binding. Although continuous co-expression of C/EBPα with OSKM increased iPS cell reprogramming efficiency for 11-fold, but only an 18-hour pulse of C/EBPα before OSKM activation resulted in a 100-fold increase in reprogramming efficiency and can reprogram 100% of the poised cells into iPS cells. Furthermore, it is indicated that transient expression of C/EBPα initiates an epithelial-mesenchymal transition (EMT), but subsequent OSKM overexpression switched EMT off and then, MET proceeds efficient iPSC generation. Although the C/EBPα technique can enhance reprogramming of B cells with 100% efficiency, however, this approach is indicated cell type specific and inoperative in pluripotent reprogramming of MEFs [53]. Thus, further investigations are needed to test the applicability of this factor on enhancing reprogramming efficiency in different cell types.

### 1.9. The Privileged Cell State

Distinct cell types have different capacities for direct reprogramming based on the tissues and species from which they are derived as well as *in vitro* environmental conditions [147]. Surprisingly, a subset of cells has recently been identified in granulocyte monocyte progenitors (GMPs), highly competent for nonstochastic reprogramming into iPS cells. Guo and colleagues named these cells as “*privileged*” cells for pluripotent reprogramming. They demonstrated that privileged cells have an ultrafast cell cycle (∼8 hr) and can be synchronously reprogrammed with a short latency [48].

It is demonstrated that there is a direct relation between cell cycle rate and reprogramming efficiency [93]. Therefore, speeding cell cycle up could induce the emergence of privileged cells [48]. Conversely, Xu *et al.* indicated that hyperproliferation might has negative effect on reprogramming efficiency [148].

Interestingly, Guo group produced a small population (1%–8%) of ultrafast cycling MEFs from normal MEFs by transient overexpression of Yamanaka factors for a limited time window (6 days) [48] before establishment of pluripotency similar to that of the cell activation phase in the cell activation and signaling directed (CASD) reprogramming approach (will be discussed in section 5-1) [62-64, 149]. However, there is evidence that indicates transient acquisition of pluripotency during the short-term OSKM treatment [150] and challenges the privileged state [151]. Interestingly, Guo *et al.* showed pluripotent reprogramming of the ultrafast MEFs more efficient (∼99.7%) than that of the normal MEFs [48]. These cells also were named as “lucky” cells, because of their commitment toward an iPSC fate [151]. It is indicated that these lucky cells only assumed a fast and accelerated cell cycle, that is a feature of pluripotent cells and nonstochasticity is unlikely to occur. Accordingly, the term ‘privileged somatic cells’ for partially pluripotent committed MEFs seems controversial [151].

Interestingly, it is revealed that depletion of p53 and p57 by a cell-cycle acceleration mechanism controls emergence of ultrafast cycling cells and increases reprogramming efficiency of hematopoietic stem and progenitor cells (HSPCs) as well as Lin−/c-Kit^+^/Sca^+^ (LKS) cells [48]. In line with the induction of “privileged state” (if be verified by experts), different factors might facilitate the emergence of this state, such as factors and methods that discussed in early section of this review (e.g. up/down (?) regulation of Mbd3 [49, 50, 54], down-regulation of p57, p16^Ink4a^/p19^Arf^, p21^Cip1^ and p53 [48, 52, 87, 95, 96, 98-100, 102, 105]), fast cell cycle kinetic, inherent populational homogeneity [48], endogenous expression levels of OSKM similar to that of stem/progenitor cells, and the level of cellular differentiation.

Collectively, these findings suggest that by identification, stimulation and isolation of ultrafast cycling cells from a known cell line (e.g. stem/progenitor cells) iPSC generation could be achieved in a deterministic level. This finding represents an intrinsic capacity of such these cells for repopulation in specific situations (e.g. injuries). Ultimately, this inherent capacity might be stimulated and induced *in vivo* as a novel strategy for induction of regeneration.

### 1.10. Combinatorial Modulation of Barriers and Enhancers

In the previous sections, we learned about different barriers and enhancers of reprogramming. In this study removal of barriers is considered as enhancing strategies. Thus, simultaneous removal of barriers and activation or administration of enhancers would have a cumulative and maximal effect on improving reprogramming efficiency and kinetics. However, this notion could be effective in the presence of synergism and in the absence of unexpected and neutralizing interactions. In some cases, it is indicated that particular pathways that act as barriers to reprogramming have interactions, and subsequently combinatorial effects to oppose the reprogramming process [118]. For instance, the clathrin-mediated endocytosis and TGF-β signaling pathways have a positive linear interaction during reprogramming that antagonizes reprogramming and subsequently decreases the efficiency [118]. Thus, some incompatibilities may exist between enhancing methods/factors that could override the effects of each other [93]. On the other hand, inhibition of multiple barriers would have an increasing effect on improving the efficiency. For example, small molecule Pitstop 2 (endocytosis inhibitor) as well as shRNAs for ADAM29 (ADAM metallopeptidase domain 29) and ATF7IP (a chromatin regulator) can enhance reprogramming efficiency up to 15-fold in a synergistic manner [118]. Interestingly, as a proof of concept for the increased effects of combinatorial modifications, Vidal and colleagues in a recently published paper have shown that modulation of specific signaling pathways (Wnt/β-catenin, TGF-β and MAP kinase) and chromatin state (by ascorbic acid and Kdm2b) can synergistically enhance the efficiency of reprogramming to a deterministic level, in a nonstochastic manner and with a reduced kinetics [71]. Their combinatorial method has caused reaching one of the highest efficiencies that is reported for highly efficient (80-100%) pluripotent reprogramming [71]. Therefore, this finding is the best proof of principle that a combinatorial method of removing barriers and activation of genetic and epigenetic enhancers can progress pluripotent reprogramming with high efficiencies and with reduced kinetics. Moreover, chemical substitutes for OSKM may help making this efficient approach a safer process (advanced combinatorial method in figure 2).

Consequently, by utilizing different findings that reported for highly efficient induction of pluripotency [48, 50, 51, 53, 75, 76, 99, 152], as discussed above, all the various methods that were developed for iPS cell reprogramming could be improved in terms of efficiency, quality and feasibility by combinatorial modulation of barriers and enhancers (advanced method in figure 2).

## 3. Chemical Approaches to Reprogramming Cell Fate

To date, numerous small molecules have been introduced in the field of stem cell biology capable of governing self-renewal, reprogramming, transdifferentiation and regeneration. These chemical compounds are versatile tools for cell fate conversion toward desired outcomes [41-43]. Small molecules have special advantages such as temporal and flexible regulation of signaling pathways, long half-life, diversity on structure and function and also their cost-effectiveness. Moreover, chemical approaches have superiority over arduous traditional genetic techniques in several aspects. For instance, rapid, transient and reversible effects in activation and inhibition of functions of specific proteins are of their profits [41, 42]. Additionally, their effects could be adjusted by fine tuning concentrations and combinations of different small molecules [41, 42]. In line with this topic, more recently, Jaenisch group report identification of a unique and chemically defined culture condition to capture a distinct naïve pluripotent state in human ESCs similar to that of the mouse ESCs using five small molecules [153]. This finding shows that chemical approaches are changing the concept of pluripotency in human. Therefore, chemicals are powerful tools in cell fate conversion and study of stem cell and chemical biology *in vitro* and *in vivo* [41, 42]. To date, chemicals have been used to improve efficiency and kinetics of traditional reprogramming method, and moreover, to substitute reprogramming factors.

To enhancing reprogramming efficiency using small molecules, it has been shown that chemical inhibitors of MEK (PD0325901) and TGFβ (SB431542) pathways can facilitate MET and improve the efficiency and kinetics of human OSKM reprogramming up to 100-fold and moreover, up to 200-fold, if small molecule thiazovivin (a ROCK inhibitor) further supplemented [86]. It is demonstrated that small molecule ascorbic acid can participate in silencing of somatic program, and has positive effects on improving quality and efficiency of reprogramming [71, 124, 130, 154, 155].

In keeping with the application of chemicals instead of reprogramming factors in pluripotent reprogramming, it has been shown that a combination of two small molecules, BIX-01294 and BayK8644, can substitute c-Myc and Sox2 [156]. Moreover, small molecule EPZ004777 can increase efficiency of mouse and human reprogramming three-to fourfold and also can substitute c-*Myc* and *Klf4* by inhibiting catalytic activity of DOT1L, a H3K79 methyltransferase and upregulation of *Nanog* and *Lin28* [125]. Compound δ is another novel synthetic small molecule that could rapidly induce several pluripotency genes in MEFs. This compound could significantly induce expression of endogenous *Oct4* and *Nanog* during cellular reprogramming [157]. Moreover, Ichida and colleagues showed that small molecule RepSox (E-616452) can replace Sox2 [85]. In addition, Zhu and colleagues demonstrated that ectopic expression of *Oct4* alone with a defined small molecule cocktail substituting for three reprogramming factors (Sox2, Klf4, and c-Myc) can induce reprogramming of several human primary somatic cell types into iPS cells [158]. However, this iPSC reprogramming procedure using small molecules and only forced expression of *Oct4* takes at least 5 weeks [158] that represents slow kinetics of this approach. Likewise, Deng and colleagues reported the induction of pluripotency in mouse fibroblasts using forced expression of only Oct4 and small molecules substituting for Sox2, Klf4 and Myc [159]. In addition to mentioned compounds, small molecules OAC1/2/3 (Oct4-activating compound 1/2/3) can activate *Oct4* and *Nanog* promoters, enhance reprogramming efficiency and dynamics and shorten the process [160]. Indeed, such these compounds potentially could be convenient substitutes for iPSC transcription factors. Notably, most compounds that can substitute reprogramming transcription factors are also capable of improving the conventional reprogramming process.

Surprisingly, Deng group in another work demonstrated that forced expression of pluripotency master genes is dispensable in iPS cell reprogramming [32]. They reprogrammed mouse somatic cells into pluripotent cells using a combination of 7 small molecules of VC6TFZ (CHIR99021, 616452, Forskolin, DZNep, TTNPB and valproic acid) plus C and P (PD0325901) in a stepwise procedure and with a frequency up to 0.2%. The produced cells were named as chemically induced pluripotent stem cells (CiPSCs) [32]. However, the main disadvantage is that chemical reprogramming takes about 40 days in mouse and it may need an even longer period for the generation of human CiPSCs [32].

Although, chemical approach is one step toward a safe reprogramming method but the efficiency and kinetics of the process is very low. Suggestively, this approaches might be improved in terms of efficiency and kinetics by identification of new small molecules substituting enhancing methods discussed formerly, (e.g. inhibition/overexpression (?) of Mbd3 [49, 50, 54], overexpression of C/EBPα [53] and down-regulation of p57, p16^Ink4a^/p19^Arf^, p21^Cip1^ and p53 [48, 52, 87, 95, 98-100, 102, 105, 106]), to develop a rapid CiPSCs production strategy. These findings demonstrate that somatic cells can be reprogrammed into pluripotent cells by using only small molecules without exogenous “master genes” and likely more efficient than ever in combination with enhancing methods. Ultimately, application of chemical modulators could be regarded as a promising strategy for development of new drugs intended for targeting endogenous (stem) cells and induced regeneration [41-43].

## 4. A Model for Simple Reprogramming of Stem/Progenitor Cells into iPSCs

In the previous sections we learned about developed techniques for enhancing iPSC reprogramming efficiency, how removing reprogramming roadblocks can facilitate and accelerate OSKM reprogramming function and how it is possible to iPSC transcription factors be substituted with specific small molecules in chemical reprogramming approaches. It has long been known that mesenchymal stem cells (MSCs) or stem/progenitor cells express certain pluripotency regulators in addition to mesenchymal markers. For instance, MSCs from human bone marrow [161, 162] adipose tissue, heart, dermis [161] and dental pulp [163, 164] expresses some key pluripotency genes (e.g.*Oct4*, *Nanog*, *Sox2*). Moreover, there are similar profiles between classes of OCT4 target genes and similar regulatory circuitries for OCT4 in hESCs and human bone marrow mesenchymal stem cells (hBMSCs) [162]. Interestingly, it has been shown that Oct4 and Sox2 (OS) can reprogram cord blood-derived CD133^+^ stem cells into iPSCs, whereas this process did not succeed in the generation of iPS cells from differentiated keratinocytes and fibroblasts [165]. Furthermore, Kim and colleagues showed that Oct4 alone is sufficient for the generation of iPSCs from mouse and human neural stem cells (NSCs), which endogenously express Sox2, c-Myc, Klf4 and SSEA1 [166, 167]. In addition to MSCs as potential resources for efficient iPSC production, it is suggested that the cells derived from visual (choriocapillaris endothelium) and immune systems might serve as competent resources for efficient reprogramming [103].

Interestingly, Eminli *et al.* showed that hematopoietic stem and progenitor cells can be reprogrammed 300 times more efficient than terminally differentiated cells [168]. Vidal and colleagues have recently demonstrated that specific progenitor cells have simpler requirements than fibroblasts for highly efficient and synchronous reprogramming that is representative of a “primed” state in stem/progenitor cells for efficient (∼95%) acquisition of pluripotency in a defined condition. This property may be due to some intrinsic features of progenitors (e.g. GMPs), including expression of stemness related genes, permissible chromatin state, a decreased level of barriers (e.g. TGF-β and MAP kinase pathways) and increased levels of genetic and epigenetic facilitators (e.g. KDM2B) that favor reprogramming [71]. Indeed, differences between embryonic and mesenchymal stem cells may originate from specific barriers (e.g. the Mbd3/NuRD chromatin repressive complex, p53 and etc.) which, in a fine tuning process, regulate gene expression levels desired for maintenance of pluri- or multipotency. By removing such these barriers of the reprogramming process and reaching a high efficiency in iPS cell production, it may be possible to convert stem/progenitor cells, which have a reservoir of reprogramming transcription factors, into iPS cells without exogenous expression of pluripotency factors. Collectively, inhibition of reprogramming barriers using small interfering RNAs (siRNAs)/small molecules and activation of reprogramming factors by chemical substitutes offer ideas for a simple method of iPSC production from MSCs *in vitro*, and additionally, provide promising hopes for stimulation of tissue specific progenitor cells *in situ* to proliferate in order to regenerative purposes. Interestingly, having approximately fast cell cycle kinetics and endogenous expression of some reprogramming factors confer a unique property to stem/progenitor cells relative to differentiated cells to be potentially elite cells poised to become iPS cells even without exogenous reprogramming factors by the combination of methods discussed in previous sections, including removing roadblocks and application of enhancers.

## 5. Cell Fate Conversion

Generation of specialized cells of various tissues attracted considerable attention as a promising hope for medical purposes. Specialized cells could be generated by distinct approaches that differ based on the starting cells and their cellular and molecular mechanisms. “Differentiation” process allows for derivation of various cell types from stem/progenitor cells by appropriate developmental cues. Notably, one potential application of pluripotent reprogramming is that somatic cell-derived iPS cells can undergo differentiation to produce desired cell types. Interestingly, somatic cells also can be converted into different specialized cells using direct lineage conversion or direct reprogramming and/or transdifferentiation in the absence of a pluripotent state. These methods utilize forced expression of lineage-instructive transcription factors as a strategy for the generation of various cell types [61, 169-173] by bypassing the multiple steps of lineage specification during development [174]. Direct reprogramming using lineage-instructive transcription factors was first done by the conversion of fibroblasts into myoblasts by overexpressing Myod [4], and subsequent studies showed the feasibility of this strategy [175-178]. Generally, there are two routes for direct lineage conversion. The first method utilizes overexpression of lineage-specifying transcription factors for direct reprogramming of cell type A into cell type B [141, 142, 179-187]. In this regard, various cells (e.g. cardiomyocytes [142, 179, 180, 188], neural cells [141, 181, 189], hepatocytes [182-185, 190]) have been directly generated from somatic cells (mostly fibroblasts) using overexpression of defined sets of transcription factors. Reportedly, there is not an intermediate state (e.g. pluripotent state or prime state) during the direct lineage conversion [61]. However, this method needs examination of many TFs and a process of elimination for identification of a key set of reprogramming factors. Furthermore, the authors showed that lower and higher expression levels and stoichiometry of master regulators could result in different fates and qualities [66, 191-193] and that the manufactured cells via direct conversion have less similarity to their *in vivo* correlates than generated ones through directed differentiation [65, 66]. The second approach, by a different mechanism, uses iPSC transcription factors (iPSC-TFs) in conjunction with appropriate conditions (soluble signals) favoring lineage specification to do such conversion through a transitory unstable state.

### 5.1. Cell-Activation and Signaling-Directed (CASD) Lineage Conversion

Ding and colleagues have demonstrated that by modifying the iPS cell reprogramming process, different cell lineages could be achieved as outcomes [62]. Based on this paradigm, starting cells (e.g. fibroblasts) transiently become “activated” by overexpression of iPSC-TFs as an early “unstable” intermediate stage and then lineage-specific soluble signals (cytokines and small molecules) redirect them toward diverse specific lineages [63] (Figure 2). They termed this approach cell-activation and signaling-directed (CASD) lineage conversion, CASD transdifferentiation or pluripotent factor-induced transdifferentiation [63, 194]. Interestingly, Yang *et al.* and Polo *et al.* have recently defined appropriate markers for a special ‘prime’ stage as Thy1^−^/SSEA1^−^ status during pluripotent reprogramming of MEFs prior to any fate commitment [103, 195]. In this cell-fate-decisive stage, the subsequent specified signaling pathways would define a specific fate in reprogramming cells [103] (Figure 2). Surprisingly, this special prime stage is in concordance with the unstable state that was reported during the CASD lineage conversion. According to this possible interpretation, in the current review, we have considered the unstable stage of the CASD transdifferentiation in mouse as the Thy1^−^/SSEA1^−^ cell state (Figure 2). Notably, Ding *et al.* speculated that transient expression of iPSC-TFs during cell activation phase erases the epigenetic identity of the examined (starting) cells [62, 64]. Interestingly, Tomaru and colleagues have recently shown that knockdown of fibroblast transcriptional regulatory networks (TRNs) in human fibroblast cells followed by a specific chemical*/*cytokine cocktail can lead to transdifferentiation of fibroblasts without expression of any transgene [196]. Suggestively, disruption of the somatic gene regulatory network may erase starting cell identity and cause an unstable or plastic state similar to the action of iPSC-TFs in the CASD paradigm and may provide a novel strategy for enhancing pluripotent reprogramming and transdifferentiation [196]. Thus, removing cell identity by selective knockdown of somatic master genes and pushing the cells in any direction by specific soluble signals offer ideas for the production of various cell types without counting on ectopic expression of virus-mediated transgenes.

Indeed, in the unstable or prime state, the starting cells lose their somatic program but do not acquire a pluripotent one [62, 64, 103]. Thus, pluripotency factor-induced plasticity is a means of dedifferentiating somatic cells and to prepare them for desired transdifferentiation [197]. Mitchell *et al*. showed that the unstable state is distinct from the state of the cells that are on the trajectory of human pluripotency acquisition (initialization, early or late stages) [197]. Whether the primed cells are “partially reprogrammed iPSCs” [127, 198-200] [124] [201] or ‘pre-iPSCs’ [90] seems unlikely. Although here we consider the state of primed cells distinct from that of partially reprogrammed iPS cells, however, it is difficult to exclude the possibility of passing through a pluripotent state and differentiation in some of the trans-differentiating cells [150, 202]. Nonetheless, further investigation is required to determine the characteristics of the primed or unstable state. Furthermore, for cell therapy purposes, it is important to generate safe progenitor or differentiated cells without residual pluripotent cells which may have teratoma-forming propensity. To this end, it is essential to avoid pluripotency state during transdifferentiation, and the safety of produced cells should be thoroughly evaluated. However, this problem is more highlight in the process of differentiation of patient-derived iPSCs than direct reprogramming and the CASD approach.

It is notable that Oct4 is capable of inducing a state of plasticity with the aid of a special reprogramming medium in human fibroblasts [197, 203]. Moreover, expression of Oct4 alone was reported sufficient for several fate conversions by the CASD paradigm [63, 197, 203-205]. Interestingly, the authors showed that Oct4-induced plasticity can transcriptionally prime human fibroblasts into multiple fates by up-regulating a subset of lineage developmental genes from all three germ layers [197]. Consequently, as possible underlying mechanisms, pluripotency factors may directly activate transcription of lineage targets, have lineage specification role [73, 206-208], interaction with epigenetic facilitators [174, 209], nonphysiological binding to lineage-specific target genes [172, 210], activate early developmental genes [197] and silence somatic program during the CASD transdifferentiation. However, it is noteworthy that factor-induced plasticity may not possess a factual *in vivo* correlate [66].

Interestingly, endogenous pluripotency genes remain silenced during lineage induction, while ectopic expression of reprogramming factor(s) is essential for the CASD reprogramming approach [62, 63]. In addition, prolonged expression of reprogramming factors in the CASD paradigm changes the balance in favor of pluripotency and inhibits transdifferentiation, likely because commitment to a specific lineage initiates quite rapidly (before induction of pluripotency) [62]. It is revealed that pluripotency conditions inhibit lineage specific conversion in the CASD paradigm [62, 63, 149, 194, 205] and for optimal lineage conversion, it is essential to have an activation stage that should be inactivated before endogenous establishment of pluripotency [62]. Additionally, authors indicated that generated cells by the CASD method, directly transdifferentiated from fibroblasts and did not pass a pluripotent intermediate stage [62-64, 149, 194, 204, 205, 211-214] (Table 1). Interestingly, findings showed that the CASD lineage specification, is not only a collection of distinct steps but also a continuous procedure in which each signal influences subsequent and preceding stimulated signaling pathways [64]. Indeed, administration of lineage specification signals with iPSC reprogramming factors in the cell activation step can improve the efficiency and quality of transdifferentiation. Suggestively, activation of lineage specific transcriptional programs in the cell activation step can specifically redirect epigenome remodeling toward lineage specification before the establishment of an intermediate stage and ensures not production of pluripotent cells [64]. Contrasting these reports, it has been shown that Nanog, a marker for the final stage of pluripotent reprogramming, is reactivated during the CASD transdifferentiation [150, 202] and enhances the efficiency of the process [150]. Accordingly, Hanna and colleagues have recently reported that vast majority of differentiated cells derived from the CASD transdifferentiation system transiently pass through a pluripotent state before somatic differentiation [150]. Thus, more investigation is needed to elucidate the exact molecular characteristics of primed cells.

**Table 1.**
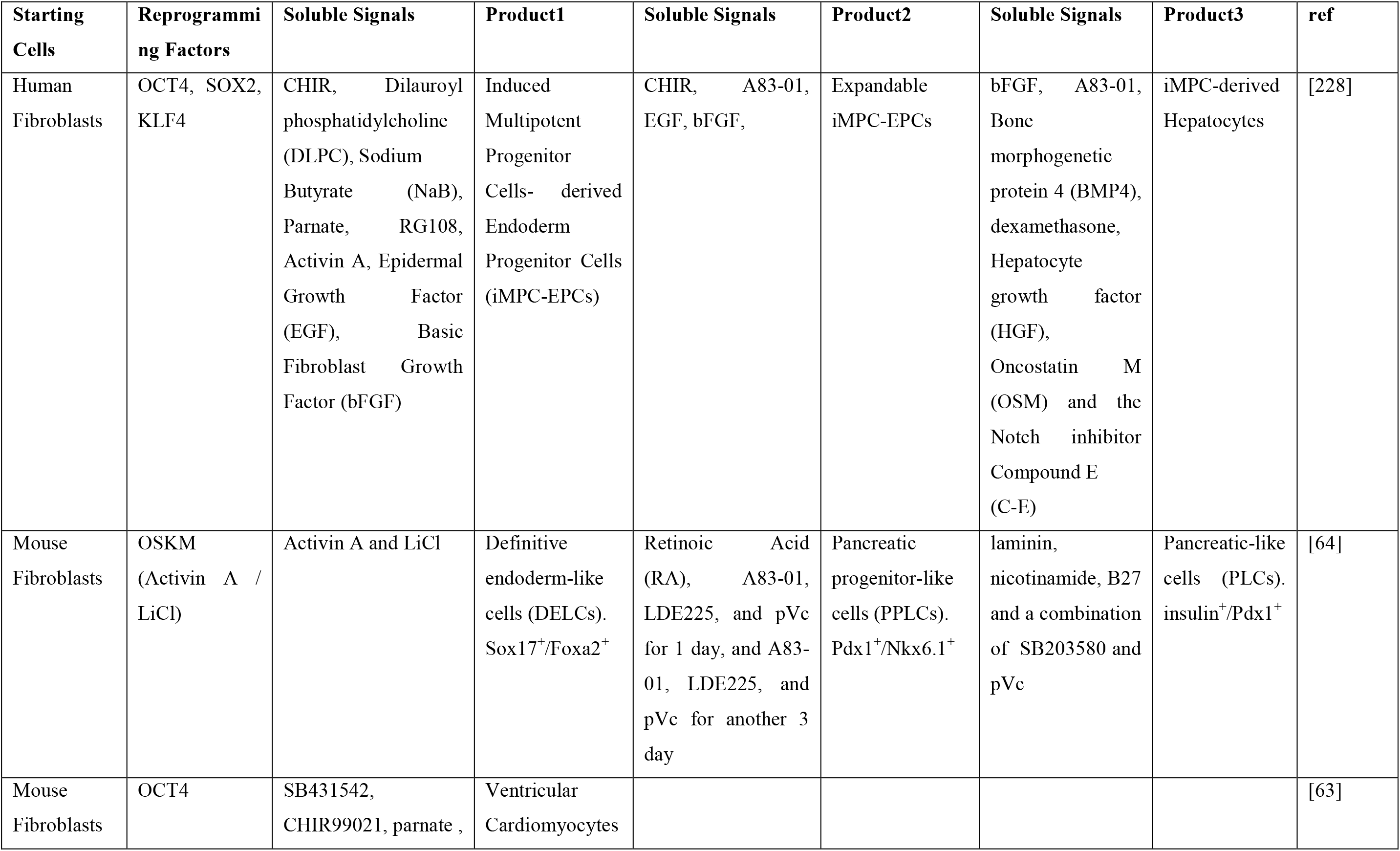
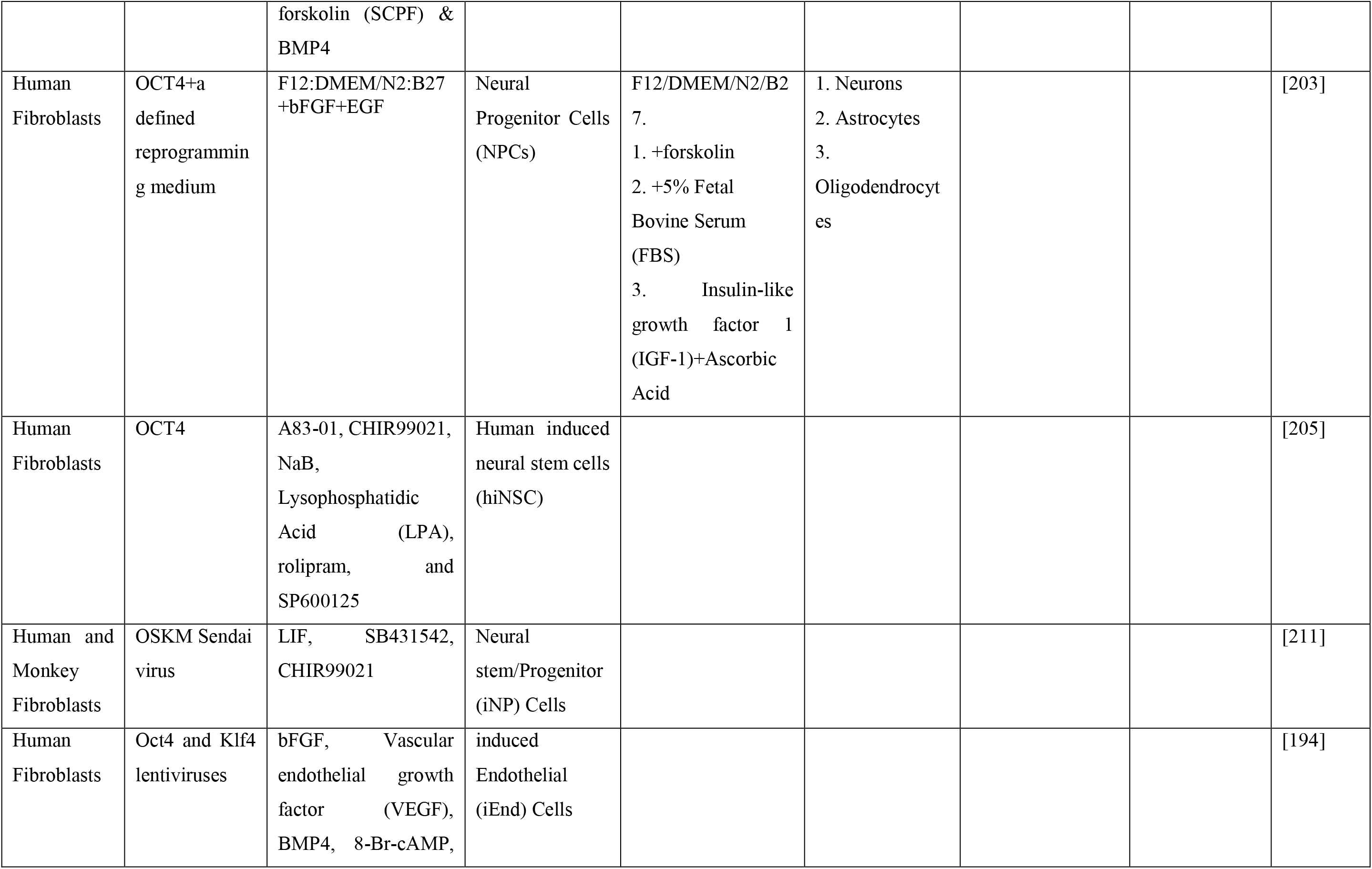
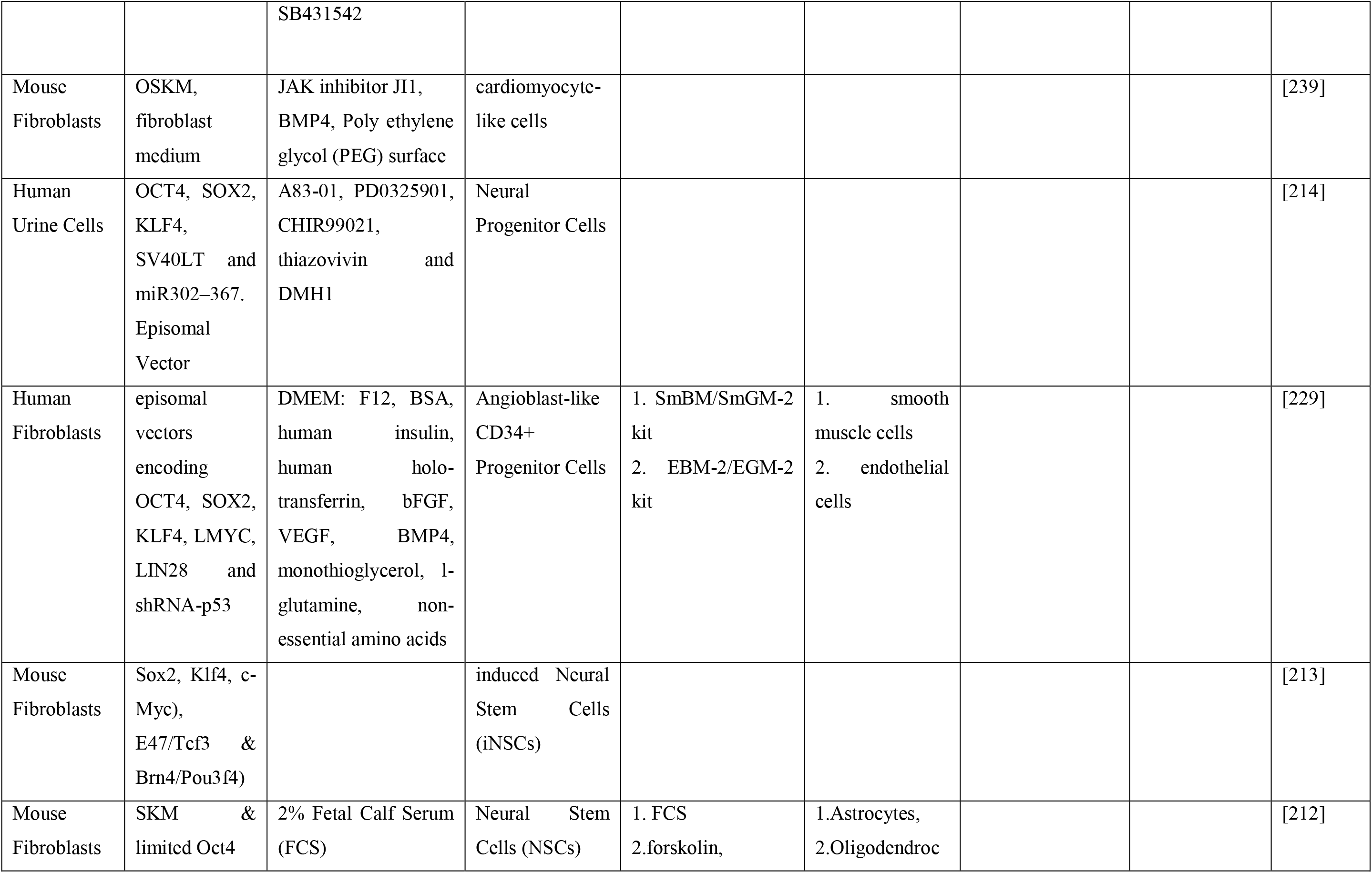
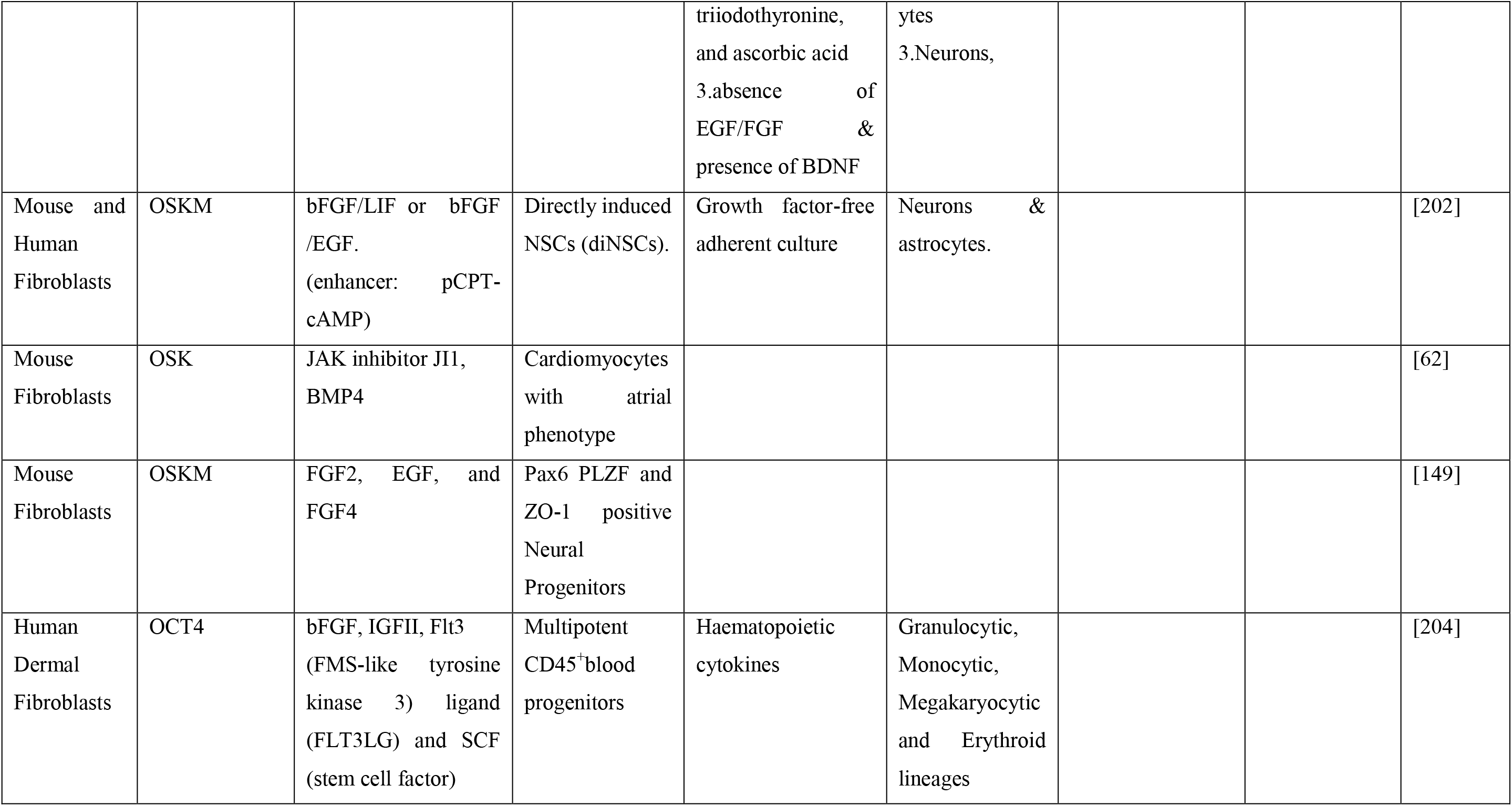
Some of the reports of the CASD lineage conversion.

Direct reprogramming by using appropriate culture conditions and forced expression of specific transcription factors responsible for progenitor state is a method for conversion of somatic cells into stem/progenitor cells. For instance, this approach has generated different progenitors, including neural stem/progenitor cells [213, 215-218], oligodendrocyte progenitor cells [219, 220], hepatic stem cells [221] nephron [222], cardiac [223], endoderm [66] and hematopoietic [224-227] progenitors. Surprisingly, the CASD lineage conversion can generate tissue-specific proliferative progenitors as well as terminally differentiated cells [64, 149, 194, 197, 204, 205, 211-213, 228, 229] (Table 1). To date, various stem/progenitor cells have been produced using this approach, including cardiac progenitor cells [62, 63], neural stem/progenitor cells [149, 197, 202, 203, 205, 211-213, 230], multipotent blood progenitors [197, 204], angioblast-like progenitors [229], expandable induced endothelial (iEnd) cells [194], expandable definitive endoderm-like cells (DELCs) [64], endoderm multipotent progenitor cells [228], and even induced EpiSCs (iEpiSCs) [231]. These intermediate cells could be considered as expandable cell sources for progenitor cell therapy purposes. Notably, the CASD transdifferentiation method has been successful in efficient generation of a variety of functional cells, such as cardiomyocytes [62, 63], pancreatic β-like cells [64], neural cells [149, 203, 205, 211-214], smooth muscle [229], hepatocytes [228], endothelial [194, 229] and blood cells [204] (Table 1).

In addition, the CASD approach can transdifferentiate derivatives of one germ layer into another that is a unique feature of this approach, for instance, it has been successful in direct conversion of non-endoderm fibroblasts into endoderm-β cell lineage [64]. Therefore, the CASD lineage conversion is a versatile approach for the generation of a variety of mature cell types and lineage-restricted stem/progenitor cells (Table 1). Contrary to what was previously thought, there is evidence that show mature assumed products of direct conversion are not terminally differentiated cells and are stem/progenitor intermediates [66]. Thus, it could be supposed that direct conversion can produce progenitor populations similar to the CASD paradigm but with fewer qualities [66].

Notably, direct reprogramming and pluripotency factor-induced transdifferentiation can be efficiently completed within a short period, as compared to the long and indirect method of reprogramming to iPS cells and subsequent differentiation into desired cell types [149]. Moreover, these methods allow for the robust generation of relevant cell types using donor/patient-derived somatic cells for cell fate conversion studies, modeling diseases and drug screening. Interestingly, CellNet analysis showed that the CASD cell fate conversion strategy gives very high target cell/tissue classification to its products than direct conversion, possibly, due to the powerful action of pluripotency factors in silencing somatic program [65] and potent effects of soluble signals in establishment of target fate.

On the other hand, as a difficulty, the CASD lineage conversion needs a fine-tuned collection of cytokines and small molecules specific for each cell types that should be administered in limited and appropriate time windows during this experimentally stepwise process (Table 1). In addition, pluripotency factors might cause stress and genomic aberrations during this transdifferentiation method [232].

In the previous sections, we address the advancements in induction of highly efficient reprogramming to pluripotency and also integration-free methods (e.g. chemical method) in reprogramming and transdifferentiation. Regarding the role of iPSC reprogramming method in the CASD paradigm, it could be suggested that this lineage conversion approach might be accomplished more efficient and without transcription factors and genetic manipulation by aforementioned techniques as a new powerful method for the production of tissue specific stem/progenitor cells and also various differentiated cell types *in vitro* and more importantly *in vivo* in order to induction of regeneration in defective and injured organs (Figure 2).

### 5.2. Native, Aberrant, and Target Gene Regulatory Networks

Thousands of genes precisely express and work together in gene regulatory networks (GRNs) to warrant the current cell’s function, steady-state, survival and its transcriptional responses to environment, disease, and age. In addition, specific GRNs are responsible for and determine the cell identity [233-235]. Although it has been neglected, but to engineer a given cell type to another fate, silencing or disruption of the native identity-associated GRNs seems essential because they can hinder the conversion process and robust establishment of the desired fate. Therefore, native/somatic GRNs that characterize a particular cell type could be considered as barriers of direct reprogramming strategies. Indeed, somatic transcriptional programs act as safeguard mechanisms in fully differentiated somatic cells by protecting the cells from aberrant transformations [196]. In this regard, Morris *et al.* showed that residual expression of native master regulators in the converted cells can repress target fate specification [66]. Interestingly, Tomaru and colleagues showed that combinatorial depletion of four fibroblastic master genes (OSR1, PRRX1, LHX9 and TWIST2) can disrupt somatic transcriptional regulatory network (TRN) in human fibroblast cells and induce a plastic state. They analyzed FANTOM5 data and Illumina microarray data to identify fibroblast-enriched transcription factors and systematic siRNA knockdown of selected transcription factors to find the minimal influential set. Moreover, they showed that ectopic expression of the four fibroblastic master genes can impede adipogenic differentiation of MSCs. [196]. Furthermore, knockdown of B cell regulators (i.e. Pou2af1 and Ebf1) during macrophage conversion of B cells can considerably improve direct reprogramming robustness at least in part through extinguish of native GRNs [66]. Thus, native identity-associated gene or transcriptional regulatory networks seem potent barriers of pluripotent reprogramming and transdifferentiation. In the case of iPSC production, although OSKM can effectively silence fibroblast GRN in iPSCs [65], however, fibroblast program appears a barrier during the reprogramming process.

Suggestively, by transient disruption of the native GRNs and subsequently unlocking the cells from somatic program diverse transformations may be possible [196]. Moreover, identification of master genes responsible for the native/somatic state in distinct cell-types and their knockdown would seem a new strategy for enhancing efficiency and fidelity of direct reprogramming to both pluripotent and mature cells. To this end, CellNet as a powerful network biology platform can identify master regulators of the native state and compare gene regulatory networks of engineered cells with their *in vivo* correlates [66]. Another issue during reprogramming is that the installation of new identity-associated GRNs in the presence of native GRNs would lead to a “confused or plastic state” but not a mature fate in the converted cells. For example, in reprogramming of B cells to macrophages [236], and fibroblasts to induced hepatocytes (iHeps) [182, 183], CellNet showed the presence of both native and target GRNs and establishment of a progenitor state instead of the intended somatic program [66]. Moreover, Cahan *et al.* showed the establishment of aberrant GRNs in products of every reprogramming strategy at least in part, due to off-target effects of the reprogramming factors and then partial establishment of alternate fates [65]. Thus, of the main molecular deficiencies of directly reprogrammed cell populations are the incomplete establishment of target GRN, persistence of native GRN, and possible existence of unintended GRNs which result in hybrid cell types [65, 66]. In addition to these problems, low network influence score of a special set of transcription factors could adversely affect the degree of target fate specification [65]. Interestingly, confirming other studies [237, 238], CellNet has revealed more fidelity of pluripotent reprogramming compared to other types of fate conversion in terms of faithful establishment of target GRNs (i.e. ESC GRN) [65]. Regarding these concerns, Morris and colleagues have challenged existing conversion strategies that could not robustly specify a defined cell fate and that engineered cells manifest minimal target fate identity [66]. In order to unraveling these problems, CellNet as a novel computational network biology platform by providing a metric of cell identity and evaluating the fidelity of cell fate conversions can prioritize the best candidates for further interventions to assist refining engineering strategies and faithful manufacture of progenitors, as well as fully differentiated cell types [66]. However, some limitations of this approach such as the inability to distinguish cell subtypes and cellular heterogeneity remain to be solved in future studies [174].

## Discussion

The current review describes recent findings in enhancement of reprogramming efficiency, chemical based reprogramming strategy in iPS cell production as one of the non-integrating techniques and the cell activation and signaling directed transdifferentiation approach. Prospectively, this study tried to explain how small molecules could be applied in the generation of iPSCs and specialized cells without application of transgenes.

The pluripotent reprogramming is an inefficient process because of various defined and unidentified barriers. In addition to the intrinsic barriers of reprogramming, the role of environmental conditions is undeniable during reprogramming. For instance, fetal bovine serum (FBS) arrests reprogramming at an intermediate stage by maintaining somatic cell program and inhibiting the activation of pluripotency genes [124]. In this regard, fine-tuning of the components of reprogramming medium could provide a powerful tool for adjusting the reprogramming rate and efficiency. To this end, small molecules are the best alternatives for defining and preparing the best optimized reprogramming milieu.

Several methods have been used for enhancing reprogramming efficiency [31, 48, 50-54, 56, 83, 87, 95-106] (Figure 2). These methods include application of inhibitors (e.g. si/shRNAs and small molecules) for such barriers as p53, p21, p57, p16^Ink4a^/p19^Arf^, *INK4a*/*ARF* locus and Bright [48, 52, 74, 96, 97], overexpression of some enhancing genes such as FOXH1 [76], C/EBP alpha [53], UTF1 [96], Glis1 [75] and INO80 [56] and also administration of some cytokines (e.g. CCL2 [55]) and small molecules (Figure 2). In the current paper, all these methods/factors were surveyed as enhancing methods/factors to introduce enhancers of reprogramming efficiency and kinetics. However, the best strategy is to avoid any genetic manipulation and over expression of enhancing genes as possible, and instead more application of small molecules.

Among the reviewed roadblocks and enhancers, some of them are more interesting and significant [48, 50, 53, 54, 196]. The most controversial barrier/enhancer of reprogramming is Mbd3 [49, 50, 54] that extensively discussed in previous sections. Another interesting and potent enhancer is C/EBPα. However, the C/EBPα is a cell type specific factor and applicable only in B cells and failed to enhances reprogramming efficiency of other cell types (e.g. fibroblasts) [53]. Therefore, identifying the implications for the applicability of this factor in pluripotent reprogramming of other somatic cells and also exploring a similar factor with a global function in other cell types with a same action to that of B cells will be interesting.

An interesting attempt on efficient reprogramming was recently reported by Guo and colleagues [48], although there is skepticism about some definitions [151]. Excitingly, they identify a special cell state known as “privileged” cell state that is highly disposed for nonstochastic and efficient reprogramming. The privileged state is a situation that both exists naturally and could be gained by alternative means as a dynamic cell state [48]. Surprisingly, ‘acquired privilege’ could be induced by transient overexpression of Yamanaka factors or specific cytokines in MEFs and LKS cells, respectively. However, distinct cell types might need different induction methods [48, 71]. Nevertheless, reprogramming efficiency and its latency varied based on the cell line and the somatic or acquired types of privileged state [48].

One of the highest efficiencies (∼100%) for pluripotent reprogramming has recently reported by Vidal and colleagues. They showed that combinatorial modulation of barrier/enhancer signaling pathways and chromatin modifiers can strongly facilitate reprogramming in a synchronous and homogenous manner [71]. Remarkably, their finding is the best proof that by a controlled combinatorial modulation of different barriers and enhancers, a highly efficient reprogramming can be achieved easily. A collection of barriers and enhancers displayed in figure 2 and advanced method explains the combined application of enhancing strategies, integration-free OSKM and OSKM chemical substitutes to introduce a highly efficient and safer reprogramming method.

According to the findings reviewed here, only depletion of barriers would be a very fast, inexpensive, feasible and accessible method for reprogramming of MSCs or stem/progenitor cells into iPS cells. However, this is an interesting notion worthy of investigation. In the final section, the CASD paradigm is reviewed as a new transdifferentiation method that utilizes primed cells at the early reprogramming steps to induce fibroblasts into a defined lineage using appropriate inductive signaling conditions. Moreover, this approach could be accomplished by non-integrating methods for a broad lineage spectrum [149].

In addition to abovementioned barriers, native transcriptional or gene regulatory networks (TRNs/GRNs) would seem to be potent barriers of pluripotent reprogramming and transdifferentiation. Here we reviewed different barriers of reprogramming, but all the barriers could be considered as subsets of somatic/native GRNs, indeed, they act inside the native GRNs. Accordingly, transient disruption of native TRN and consequently, unlocking the cells from the somatic program may provide a novel strategy for enhancing pluripotent reprogramming and together with a specific chemical*/*cytokine cocktail may be a convenient method for future transdifferentiation approaches without the need of ectopic expression of transgenes [196].

Suggestively, in tissues lacking sufficient progenitors to regenerate, induction of a progenitor-like state in resident cells and then differentiation of these induced progenitors would be a promising hope for induction of regeneration *in situ.* This method could possibly be assumed premier than direct reprogramming of a limited number of terminally differentiated cells to desired differentiated cells, (e.g. *in vivo* induced cardiomyocytes [240]). For instance, hepatic periportal oval cells which have a progenitor like state are highly competent for reprogramming into functional pancreatic β-like cells for long term, in contrast to terminally differentiated hepatocytes which cannot maintain β-cell function in the same conditions [171]. Interestingly, direct reprogramming and the CASD lineage conversion are capable of generating tissue specific progenitors in addition to functional mature cells. Although tissue progenitor identity of engineered cells may be ideal for therapeutic applications, but more robust cell fates must be engineered for applications of drug testing and disease modeling [66]. Therefore, these powerful and versatile approaches are potentially convenient alternatives for research studies, *in vivo* reprogramming and induced regeneration.

Nowadays, chemical compounds are considered as convenient tools for regulation of protein functions, epigenetics and signaling pathways that can influence quantity and quality of reprogramming products. Moreover, new findings showed that chemical factor-based reprogramming could be an almost safe, reliable, feasible, standardized and cost-effective strategy for translation of reprogramming technology into clinical applications [41, 42, 241]. Interestingly, by fine tuning concentrations of different chemical compounds in specific cocktails, their effects could be adjusted in terms of activation and inhibition of cell signaling pathways appropriate for production of various cell types.

Here we mostly emphasize application of small molecules in reprogramming methods. Although there are limitations, however, RNA interference (RNAi) using small interfering RNAs (siRNAs) is a safe strategy for transient gene silencing for *in vitro* and *in vivo* applications [242-244]. Moreover, RNAi is appropriate for reprogramming purposes than the transgene expression in current reprogramming methods [196]. For instance, siRNA transfection is a convenient tool for the removal of barriers and also transient disruption of the somatic TRN in direct reprogramming strategies (e.g. instead of OSKM expression in the CASD paradigm).

Despite reviewed beneficial aspects of direct reprogramming, there are skepticism about identity and function of directly produced cells. For unraveling some identity associated problems related to native, aberrant and target GRNs in direct reprogramming strategies, CellNet [65, 66] would help for reconsidering and refining available conversion protocols for the generation of more functional and robust cell types.

The current study has intended to explain some molecular events underlying environmental, genetic and epigenetic mechanisms of pluripotent reprogramming. Noticeably, different somatic cell types have disparate requirements for efficient reprogramming and there is not an effective general strategy for safe and efficient reprogramming in all cell types and developing such techniques is desirable. The mechanistic insights that discussed here on enhancing reprogramming efficiency and application of small molecules for replacing reprogramming factors could provide powerful tools toward safe, rapid, easy and efficient reprogramming. Accordingly, utilizing the enhancing factors and chemical approaches, would theoretically make transdifferentiation techniques almost safe and highly efficient appropriate for *in vivo* regeneration studies and presumably for clinical trials. However, presence of some degree of epigenetic memory specific to the original cell type after reprogramming still remained challenging to resolve [172, 173, 245]. Although we prospectively speculate some ideas, however, the mentioned possibilities still have to be confirmed by experimental evidence.

## Conclusions and Future Perspectives

The iPS cell technology has profoundly changed the fields of stem cell and developmental biology. In addition, the discovery of iPS cells has sparked new hopes for treating genetic and degenerative diseases. This technology has experienced dramatic progress and although to a limited scale, but reached to the clinic [246]. Recent advancements in identification of different barriers/enhancers of reprogramming and also chemical substitutes for reprogramming factors allow for near safe and highly efficient generation of the iPS cells. Applying the aforementioned techniques (e.g. highly efficient chemical reprogramming) in transdifferentiation authorize the generation of various cell types in a unique scale. These progresses shed light on both personalized and regenerative medicine purposes. Collectively, the evidence that is addressed here increases the opportunity to gain greater understanding of the reprogramming process, modeling diseases and regeneration by using specific chemicals or drugs, instead of genetically manufactured products for the generation of medically-relevant cell types. We tried to collect particular evidence of reprogramming and transdifferentiation methodologies to propose a blueprint toward safe and efficient cell fate conversion in order to clinical applications of both cell-based and factor-based therapy.

## Acknowledgements

The author would like to appreciate Dr. Russell C. Addis from the University of Pennsylvania, for discussions and editing the manuscript. Thanks go to Seyed Ahmad Tabatabaei, Seyed Khalil Forouzannia, Mohammad Hossein Soltani and Mahdieh Namayandeh for their administrative supports. The author apologizes to all scientists whose excellent works could not be discussed in this review due to space restrictions.

